# Alterations in amygdala function explain how aging and loneliness impair trust learning

**DOI:** 10.64898/2026.01.14.699282

**Authors:** Ronald Sladky, Federica Riva, Claus Lamm

## Abstract

Functional relationships are essential for healthy aging and they rely on social cognition skills such as establishing and monitoring whom to trust. However, aging and loneliness can negatively affect social brain function, potentially leading to a vicious cycle. Using functional MRI and computational modeling we investigated trust learning in a sample of neurotypical older (64-84 years, n=29f/23m) compared to younger adults (20-33 years, n=31f/31m). Older participants displayed lower initial trust and less trust learning when repeatedly interacting with a trustworthy and an untrustworthy trustee. Their basolateral and central amygdala activation was lower during trust decisions, and this was associated with less optimal trust behavior. Computational modeling also revealed that a crucial learning parameter, trust prediction error precision, was decoupled from basolateral amygdala activation, and this effect was pronounced in lonely older adults. These findings indicate that differences in amygdala function and reward signaling at older age and higher loneliness could drive detrimental effects on trust learning, leading to poorer social cognition and putting individuals’ sociality and well-being at risk.

## Introduction

Engaging in positive and reliable social relationships benefits well-being not only on a hedonistic level but is also an important factor for mental and brain health, as part of a multilevel resilience and resistance network (Cohen et al., 2022). It has been argued the human brain’s goal is to optimally self-regulate internal somatic homeostasis and that embeddedness in a functional social environment constitutes an important aspect in this regard (Matthews and Tye, 2019). Failures in this socially regulated homeostasis can result in loneliness, i.e., perceived social isolation, which has been discussed as a most potent threat to survival and longevity (Bzdok and Dunbar, 2020). Indeed, a recent large-scale multi-national study in 19 countries has suggested chronic deficits in social connectedness to act as a major risk factor for end-of-life symptoms (Pivodic et al., 2024) and as a predictor for worsening of depressive symptom trajectories (Raina et al., 2021), including increased risk of suicidality (De Leo, 2022) in older people.

Successful social interactions require trust in other people but also well-calibrated mechanisms in social learning to mitigate the potential of exploitation by others. Accumulated negative lifetime experiences can impair trust behavior (Bell et al., 2019). While such experiences may have protective effects, it may also entail lost opportunities if trusting would be a safe and beneficial option. Biases in social trustworthiness have also been linked to mental health issues (Roychowdhury, 2021). In psychotic disorders, for instance, flexibility in trust behavior in response to socially relevant information is considered a critical factor for dysfunctions in social relationships and may drive progression from subclinical symptoms to full-blown psychosis (Fett et al., 2012).

In behavioral research and neuroscience, trust has traditionally been investigated using single-shot trust games that target *a priori* beliefs about the trustworthiness of other people (Berg et al., 1995). More recently, researchers, including from our group, have studied trust learning by using a repeated version of the trust game, where participants need to learn whom to trust, and whom better not to trust (Mikus et al., 2023; Rosenberger et al., 2019; Sladky et al., 2021). Computational modeling of this task alongside targeted analyses of subcortical nuclei have revealed important new insights into the neuro-computational mechanisms of trust and trust learning. In the present study, we exploited the same task and analysis approaches to extend our knowledge on how the amygdala’s and connected areas’ involvement may explain how trust changes with aging and loneliness.

To this end, we collected data in a sample of older adults (n=52, 64 to 84 years of age) to test the influence of aging and loneliness on trust learning on a behavioral (investments), computational (learning parameters), and neuronal level (fMRI BOLD response) in comparison to a sample of younger adults (n=62, 20 to 33 years of age). Behaviorally, we hypothesized that older adults would be less trusting and adaptive in their investment behavior (less differentiation between a trustworthy and an untrustworthy trustee), resulting in suboptimal monetary gains from the trust interactions. Computationally, we hypothesized this suboptimal behavior to be tied to lower learning rates and beliefs about the trustee’s volatility, which both reduce the rate at which trustworthiness beliefs are updated. Finally, since increased dopamine availability has been shown to improve trust learning (Mikus et al., 2023), and since aging and loneliness/isolation have been linked to loss of dopaminergic function, we predicted a role of dopaminergic midbrain areas in trust learning (Schuster and Lamm, 2025) and their interactions with the amygdala to underpin these differences in trust and trust learning.

## Results

### Trust behavior is lower in older adults

52 healthy, neurotypical older adults (O group, age: 64 to 84 years) and 62 younger adults (Y group, age: 20 to 33) participated in this study (data from the younger adult sample have been reported previously (Sladky et al., 2021)). The older adults sample has not been reported elsewhere, and the present study focuses on comparisons across the two age groups, their interaction with loneliness, and a refined computational modeling approach to investigate trust and trust learning. Loneliness groups were matched for group size, age range, and gender distribution. All participants had no history or current diagnosis of neurological or psychiatric disorders. They had been preselected to show either high (L+) or low levels of self-reported loneliness (L-), assessed by the UCLA Loneliness questionnaire, and to be between 20 to 35 years or older than 60 years old. Demographical data are reported in the methods section and in Table 1.

**Table 1:** Demographical data.

|  | Older Adults |  | Younger Adults |  |
| --- | --- | --- | --- | --- |
| Loneliness | L- | L+ | L- | L+ |
| n | 25 (12f/13m) | 27 (17f/10m) | 31 (15f/16m) | 31 (16f/15m) |
| Age [Years] | 70.0 ±3.5 | 70.9 ±5.5 | 23.6 ±3.2 | 24.0 ±3.0 |
| UCLA Loneliness Scale | 27.6 ±3.1 | 40.7 ±7.6 | 29.1 ±4.9 | 47.6 ±6.5 |
| Education [Years] | 13.4 ±3.1 | 12.9 ±1.9 | 15.0 ±2.6 | 14.5 ±1.8 |

While undergoing fMRI scanning, participants played investors in the repeated trust game for 20 rounds with each of two trustees (2 × 20 rounds, with trustees alternating). Each round, participants invested between 1 to 10 points, which were transformed to Euros after the experiment. To increase ecological validity, participants were briefly introduced to two age- and gender-matched confederates at the outset of the experiment, who allegedly were the two trustees in the upcoming trust game. In reality, the trustees’ back-transfer behavior was simulated, with returns following a preprogrammed response schedule of a trustworthy and an untrustworthy trustee, based on prior work (Rosenberger et al., 2019).

In brief, our analysis approach combined an orchestrated set of analyses targeting different aspects of trust learning on a behavioral, computational, neuronal, and neurocomputational level, where we used linear mixed models (LMM) for trial-by-trial data, ANOVAs for aggregated per-subject data, and general linear models (GLM) for fMRI whole-brain analyses.

Analysis of the total of trust investments across the 20 rounds, separated by trustee, and using a Linear Mixed Model (LMM formula: Investment ~ Age Group × Loneliness Group × Trustee + (1|Subject)) revealed significant main effects and interactions. Total trust was lower in older adults (β=24.24, 95% CI [7.00, 41.49], t(218)=2.77, p=0.006; Std. β=0.52, 95% CI [0.15, 0.88]), the L+ Groups (β=−18.56, 95% CI [−36.37, −0.75], t(218)=−2.05, p=0.041; Std. β=−0.40, 95% CI [−0.78, −0.02]), and when investing trust in the untrustworthy trustee (β=−52.92, 95% CI [−66.52, −39.32], t(218)=−7.67, p<.001; Std. β=−1.13, 95% CI [−1.42, −0.84]). The interaction of Age × Loneliness, irrespective of Trustee, was not significant (p=0.262) but Age × Loneliness × Trustee was (β=−32.42, 95% CI [−58.00, −6.83], t(218)=−2.50, p=0.013), motivating separate ANOVAs for the two types of trustees. This revealed no significant effects for the untrustworthy trustee (all main effects and interactions p>0.137; see Supplement for comprehensive results report). For the trustworthy trustee we found significant age effects (Y>O, p<0.001) and a trend for loneliness effects (L->L+, p=0.069). The largest investment difference was between OL+ and YL− (difference=−42.80 points, 95% CI [−64.83, −20.78], p<0.001), suggesting that the age-related lower investments in the trustworthy trustee are amplified by higher loneliness.

When analyzing only the first round (i.e., with no information yet on trust behavior of either trustee) as a measure of initial trust, older adults invested less than younger adults (first round investment (±SEM) in O: 5.17±0.33 points, compared to Y 6.43±0.31, Figure 1a). Predicting initial investment with the Age and Loneliness Groups (LMM: Initial Investment ~ Age Group × Loneliness Group + (1|subject)) revealed significantly higher initial trust investments in the Y Group (β=1.53, 95% CI [0.25, 2.80], t(222)=2.35, p=0.020; Std. β=0.58, 95% CI [0.09, 1.07]) but no loneliness-related effects (all p>0.557, Figure 1b).

**Figure 1:**
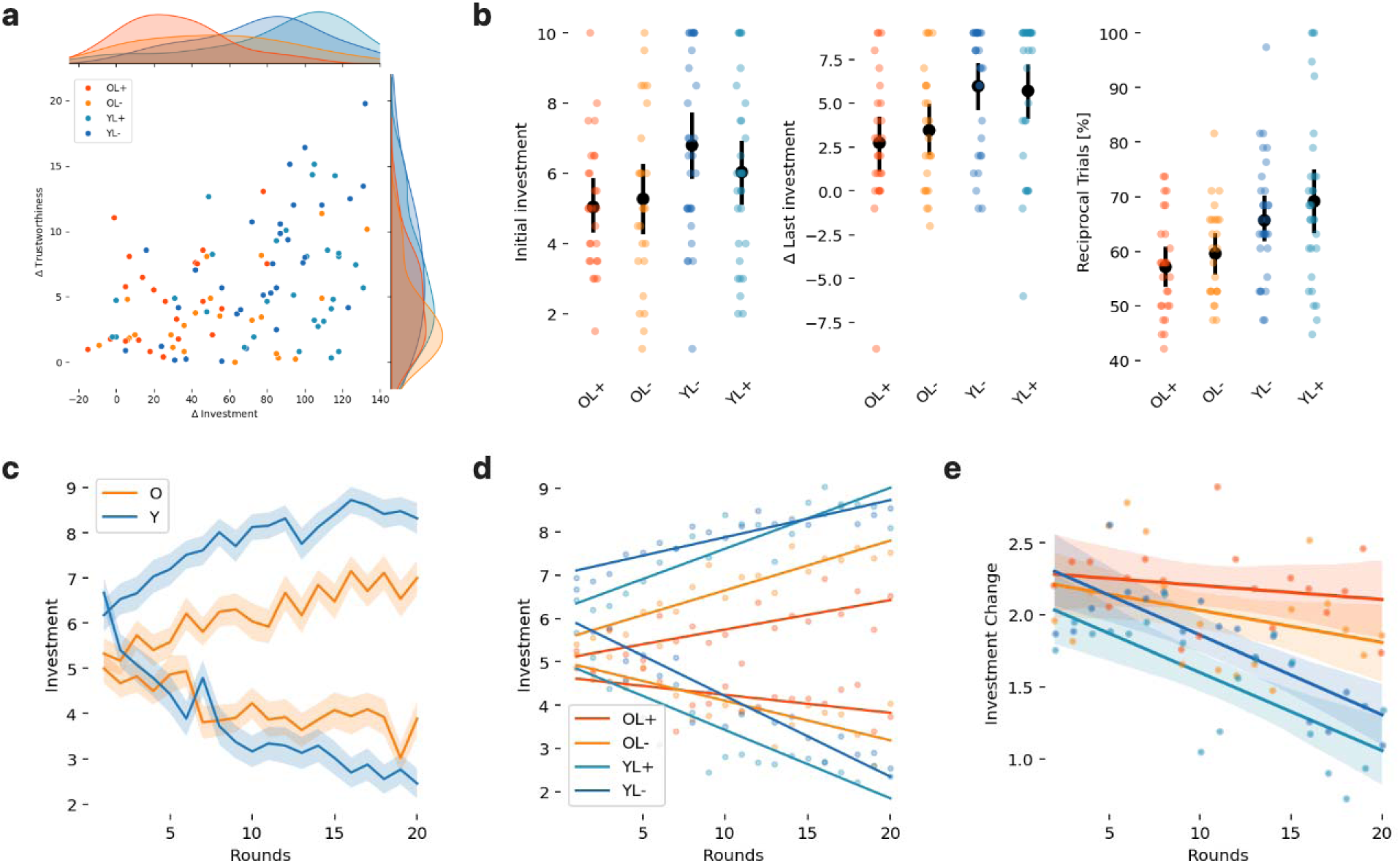
Participants played the repeated trust game in an MRI scanner, acting as investors with two simulated trustees: one trustworthy and one untrustworthy (2lJ×lJ20 rounds) **a** Compared to younger adults (Y), older adults (O) differentiated less between the trustworthy and untrustworthy trustees in both monetary investments and subjective trustworthiness ratings; even more so when they reported higher levels of loneliness (L+) (median split of the UCLA Loneliness Scale scores). **b** In older adults, we found lower first round investment (indicating lower initial trust), lower investment differences between the trustworthy and untrustworthy trustee in the last trial (indicating lower overall trust learning), and lower reciprocity (indicating lower trial-by-trial trust learning). **c** The raw investment data suggests that older adults have a flatter slope in addition to the lower initial investment. **d** Self-reported loneliness amplified the age-related differences in investment, suggesting learning to be affected negatively by higher age and higher loneliness. **e** Likewise, the same effect was observed for the investment change across rounds, indicating lower trust learning in older adults, particularly in those who were lonely (OL+). See supplementary results for full details.

In sum, the analyses of the behavioral data showed that older adults invested less trust than younger adults in the repeated trust game, both in the first round and overall, and that the latter was linked to more defensive trust investments for the trustworthy trustee, and not by differences in trust in the untrustworthy trustee. Loneliness had rather unsubstantial main effects, and did not affect initial trust; in combination with older age, however, it resulted in lower trust in the trustworthy player. These findings imply meaningful behavioral learning differences between the age and loneliness groups, to which we turn to next.

### Lower trust learning behavior in older adults across repeated interactions

First, we investigated how participants differentiated (ΔInvestments between trustees) their trust behavior and subjective trustworthiness ratings between the trustees (all rounds ΔInvestment ~ Age × Loneliness) as an indicator for their trust learning abilities. Differences in investment behavior were influenced by age (Y>O, F(1,110)=6.14, p=0.015, η^2^(partial)=0.04) and loneliness (L->L+, F(1,110)=5.17, p=0.025, η^2^(partial)=0.05) and their interaction (F(1,110)=6.24, p=0.014, η^2^(partial)=0.05). The post-hoc t-tests showed that the latter was driven by an interaction of higher loneliness and age, as the difference OL− and YL−was smaller and not significant (p=0.069), yet it was larger and significant for the lonely adults YL+>OL+ and YL+>OL− (both p<0.001). For the investment difference in the last round only (last round ΔInvestment ~ Age × Loneliness), which quantifies how the participants differentiated between the trustees after the maximum of learning options within the experiment, we found a significant main effect of age (Y>O, F(1,110)=14.64, p<.001, η^2^(partial)=0.12), but no effects of loneliness or the age × loneliness interaction (all p>0.521). Additionally, we analyzed the proportion of reciprocal investments (Reciprocal ~ Age × Loneliness), which reflects how often participants adjusted their trust behavior based on gains or losses in the previous trial, which again revealed a significant age effect (Y>O, F(1,110)=17.35, p<.001, η^2^(partial)=0.14), but no main effect of loneliness nor its interaction with age (all p>0.178, Figure 1b).

Then, we investigated changes in trust behavior across the rounds using a LMM (Investment ~ Age × Loneliness × Trustee × Round + (Round|Subject)). We observed significant effects for age (Y>O, t(4540)=2.66, p=0.008), round (t(4540)=4.44, p<.001), and Trustee × Round (β=−0.21, 95% CI [−0.26, −0.16], t(4540)=−8.23, p<.001; Std. β=−0.37, 95% CI [−0.45, −0.28]), which means that participants adapted their investment depending on the trustee across rounds (T+>T-). Furthermore, age [Y] × loneliness [L+]) × round (β=0.10, 95% CI [6.26e-03, 0.20], t(4540)=2.09, p=0.037; Std. β=0.18, 95% CI [0.01, 0.35]), loneliness [L+] × trustee [T-] × round (β=0.10, 95% CI [0.03, 0.16], t(4540)=2.78, p=0.005; Std. β=0.17, 95% CI [0.05, 0.29]), age [Y] × loneliness [L+] × trustee [T-]) × round (β=−0.12, 95% CI [−0.22, −0.03], t(4540)=−2.61, p=0.009; Std. β=−0.22, 95% CI [−0.38, −0.05]) interactions, were significant. This means that loneliness modulated learning trajectories in addition to the age and trustee effects (Figure 1c-d).

In sum, at the behavioral level, we observed significant age group differences in trustworthiness ratings, initial investments, last-round investment changes, and the proportion of reciprocal trials. Several of these effects were further modulated by self-reported loneliness, particularly total investment, investment trajectories across trials, and absolute investment change. These behavioral findings point to converging age-related differences in trust learning processes that go beyond simplistic explanations such as that older adults are inherently less trusting or more risk-averse; they also highlight a contributory, yet less consistent role of loneliness. Our next aim was, therefore, to unravel the computational processes behind these effects.

### Computational modeling reveals lower learning rates in older adults

Using a computational Bayesian belief model that implements a hierarchical Gaussian filter (HGF) (Mathys et al., 2014) and which we had previously validated using the same task (Mikus et al., 2023), we inferred the participants’ belief changes about the trustworthiness of the two trustees based on their trial-by-trial behavior (Figure 2a), using two parameters, belief volatility ω and precision weight (ψ). The model estimated the change of each participant’s precision of beliefs, using a participant- and trustee-specific parameter ω. This so-called belief volatility parameter is a measure of the participant’s flexibility (high volatility ω evinces stronger incorporation of feedback) or stability (low volatility ω prevents strong shifts of beliefs) of prior beliefs about a trustee before receiving a back transfer. Technically, a Gaussian distribution with a specific mean and precision is used to approximate the latent beliefs about a trustee’s trustworthiness. The round-by-round learning rate (ψ) on the prediction error, called precision-weight, is inversely proportional to the expected precision of beliefs. Higher values thus imply a stronger incorporation of prediction errors (occurring when back-transfers were higher or lower than expected) into updates of expectations, as expressed by the trust decisions in subsequent trials. In addition, we estimated initial trust (μ0) and choice precision (γ) based on the trust behavior. Further methodological details are documented in the methods section.

**Figure 2:**
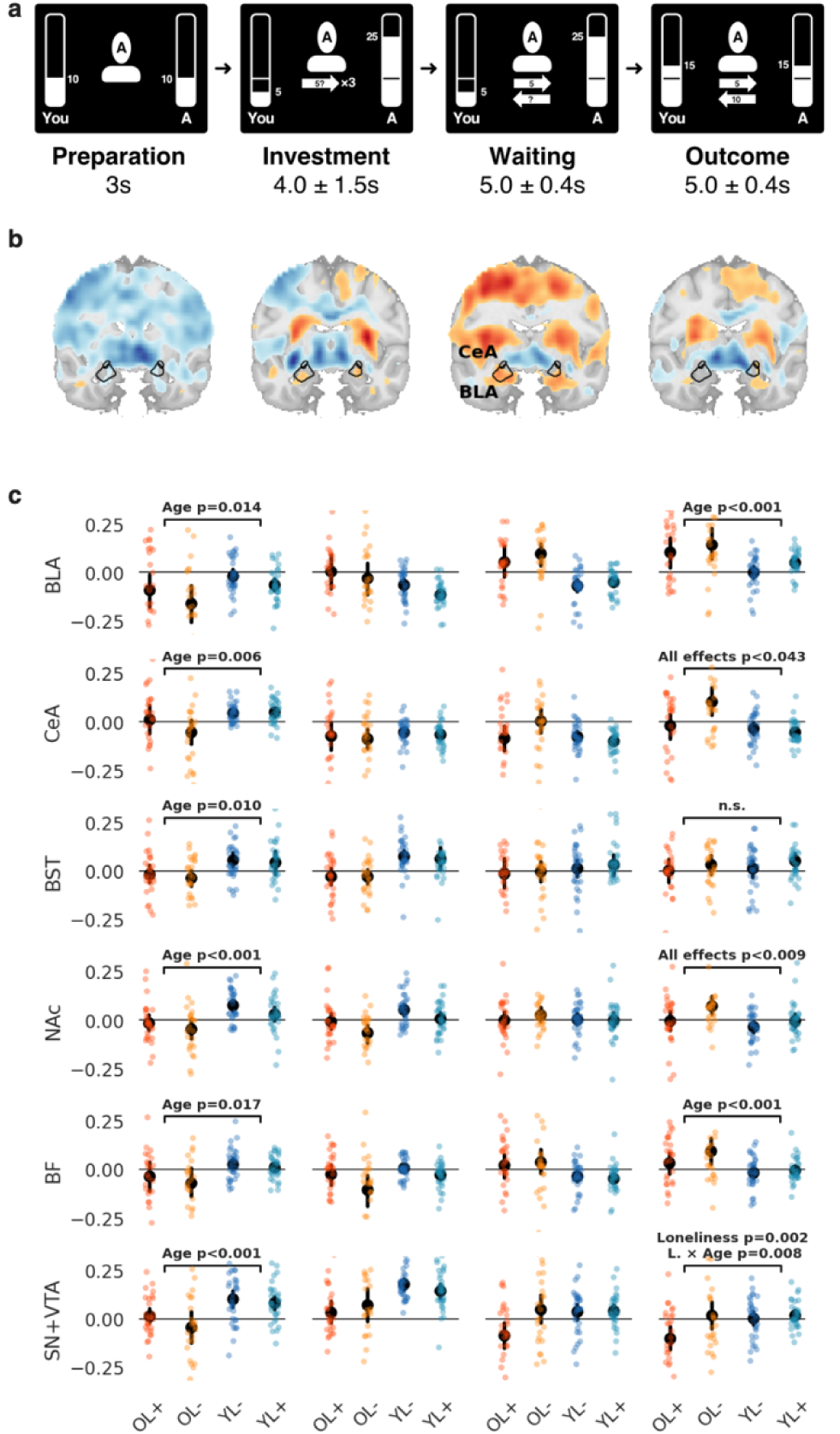
fMRI group-level results. **a** fMRI implementation of the trust game. Participants played the repeated trust game alternating with a (simulated) trustworthy and an untrustworthy trustee (2L×L20 rounds). Preparation phase: Participants were presented with the face of the trustee they played with in this round. Both received an endowment of 10 points at the outset of each round. Investment phase: Participants were asked to select an amount of 1 to 10 points to invest in the present trustee. The amount invested was tripled and added to the trustee’s account. Waiting phase: While the trustee allegedly made its decision, the participant needed to wait. Outcome phase: Finally, the trustee transferred back points to the participant, resulting in a non-negative outcome for the trustworthy (as shown in the example) and a non-positive outcome for the untrustworthy trustee. **b** Whole brain results of the older>younger adults (hot) and younger>older adults (cool). SPMs of the group contrast for both trustees combined, threshold at pL<L0.01 for display purposes. **c** During the preparation phase, which is critical for the trust decision, BOLD responses were lower in O (orange) compared to Y (blue) in the amygdala subnuclei (BLA, CeA, BST) and substantia nigra/ ventral tegmental area (SN/VTA) and basal forebrain (BF). During trust learning in the outcome phase, we found hyperactivation for O in the BLA, CeA, NAc, and BF; loneliness affected CeA, NAc, and SN+VTA.

For all parameters we calculated separate ANOVAs (parameter ~ Age Group × Loneliness Group). For initial trust (μ_0_) we found significant main effects for age (Y>O: F(1,110)=1562.63, p<.001, η^2^(partial)=0.17) and for loneliness (L->L+: F(1,110)=22.03, p<.001, η^2^(partial)=0.01), but no interaction (p=0.264). For choice precision (γ) we found no significant effects (all p>0.090).

For the learning rate (ψ) we found no significant main effects for age (p=0.114) and loneliness (p=0.265) but a significant interaction (F(1,110)=11.72, p<0.001, η^2^(partial)=0.02). Post-hoc t-tests show no significant differences between not lonely participants (YL−>OL−, p=0.387) and older participants (OL+>OL−, p=0.679) while all other differences were significant (all p<0.03) (Figure 2b). Learning rate was the lowest in the OL+ group, indicating lowest learning from prediction errors in lonely older adults. We will later analyze this mean learning rate estimated across all trials on a trial-by-trial basis and relate it to differences in BOLD activation.

For the volatility parameters (ω_mean_, ω_trustworthy_, ω_untrustworthy_) we found no significant age effect (all p>0.330). However, there were significant effects for loneliness (all p<0.003) and the interactions (all p<0.004). Post-hoc comparisons indicated that in the O group, all ω’s were lower for L+ compared to L- (all p<0.016), while no such difference existed for the Y group (all p>0.695).

As an interim summary, older adults showed the most pronounced differences in their lower initial trust estimates (μ*_0_*), consistent with the lower initial investment observable in the raw behavioral data. Choice precision did not differ between groups, but there were marked reductions in the learning rate (ψ) and beliefs about the volatility of the trustees (ω), specifically in lonely older adults. However, older adults with low loneliness resembled younger adults in this respect. Overall, this suggests that when higher age and loneliness are combined, participants show reduced learning from feedback and less flexibility in updating their beliefs about the trustees.

The modeling analyses so far aggregated across trustees and did not consider individual differences in learning performance (i.e., to which extent individual participants learned to differentiate their trust behavior between the two distinct trustees). To investigate what factors could affect the trial-by-trial learning rate (ψ), we calculated a series of linear mixed effects models. Before investigating the full model (ψ ~ Learners Group × Age Group × Loneliness Group + Trustee + (1 | subject)) we validated that ψ actually captures the expected learning goal to differentiate between the trustees (LRN+/− learners group definition based on the median split in the total investment differences (Sladky et al., 2021)). We found that the learning rate is higher for the untrustworthy trustee, more so in young, especially if lonely (see supplement and Figure 2c). Likewise, the learning rate was higher in learners and affected by age and loneliness. While non-lonely groups are comparable, ψ was highest in YL+ and lowest in OL+ (see supplement and Figure 2d).

**Figure 1:**
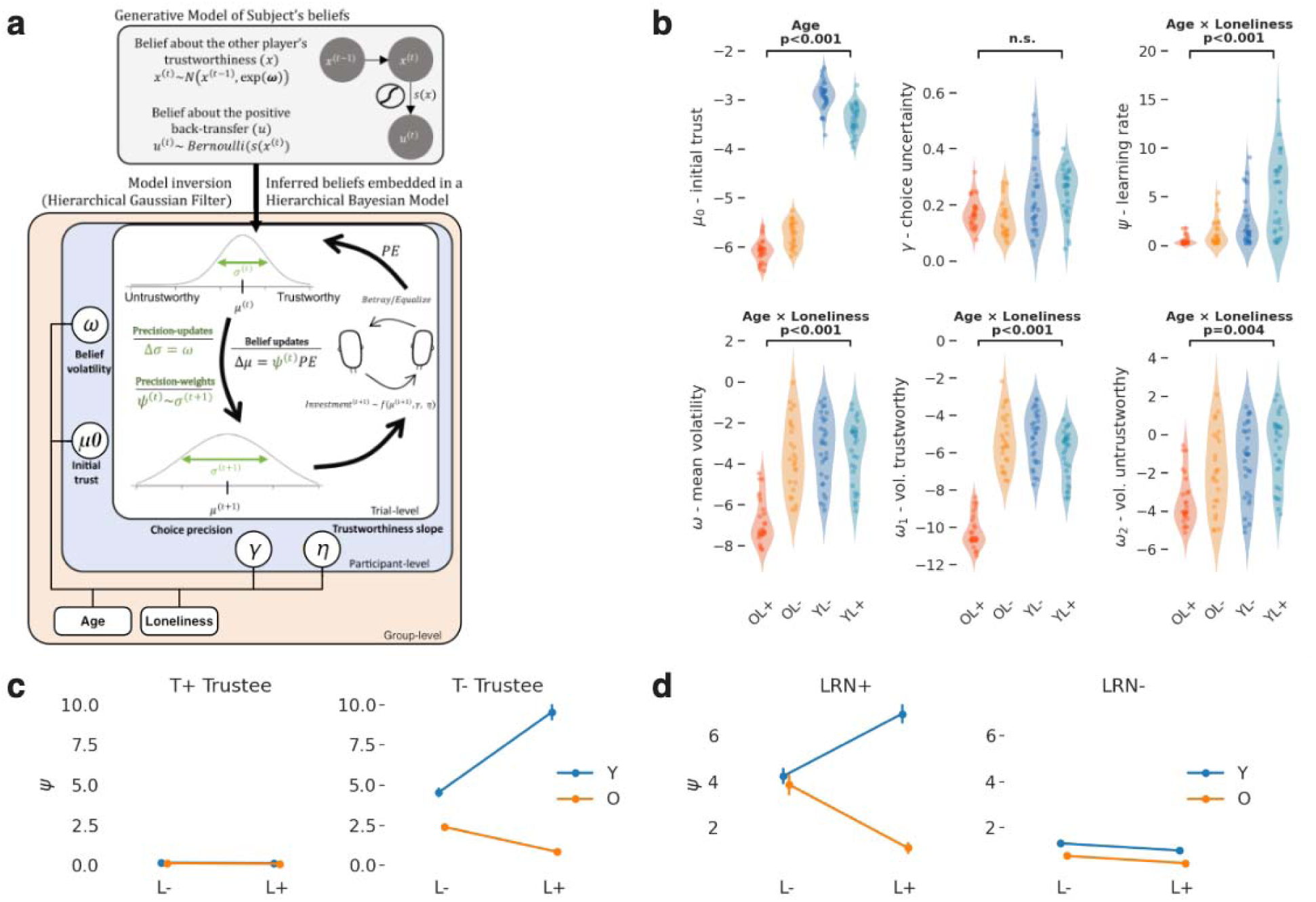
Computational modeling of participants’ behavior to infer on their beliefs about the other trustees’ trustworthiness. **a** Hierarchical Gaussian filter (HGF) model (figure adapted from (Mikus et al., 2023)). A Gaussian random walk with step size of ω was used as a generative model for trustworthiness beliefs *x* at trial *t* and inverted using the HGF, resulting in trial-by-trial estimates *N*(μ*_t_*, σ*_t_*). Belief volatility parameters ω for the trustworthy and untrustworthy trustee govern the rate of change of σ_t_ via the precision-weights ψ*_t_* that serve as dynamic learning rates. Identical to the original publication, we estimate initial trustworthiness belief per participant (μ*_0_*). The ordinal logistic link function governs how beliefs about others’ trustworthiness map to investments with two additional subject-level parameters: choice uncertainty (γ) and the slope (η). The parameter estimation is done through hierarchical Bayesian inference, where we estimate all individual and group-level parameters in one inferential step (see Mikus et al., 2023, and Methods, for further details). **b** Compared to younger adults (Y, blue) initial trust (μ*_0_*), volatility (ω) for both trustees and precision-weights (ψ) were lower in older adults (O, orange), an effect that was higher in higher self-reported loneliness (L+). There were no differences for choice uncertainty (γ). For detailed statistics, refer to the main text and Table 1. **c** The learning rate (ψ) was higher for the untrustworthy trustee, and higher in younger adults. This age-effect was amplified by loneliness. For the trustworthy trustee there were no such effects. **d** As expected, learning rates were higher in learners (LRN+); in non-learners (LRN-), on the other hand there were less age and loneliness effects.

In summary, the computational modelling analyses indicate that older adults, particularly those who feel lonely, show a lower learning rate and thus learn less from feedback (as modeled by their prediction error precision) and lower volatility beliefs (i.e., more stability and thus less flexibility in their beliefs), which both affect the way they perform learning from the outcome of positive and negative trust interactions. Next, we investigated how regional neural activity differed between the groups and how this may explain the behavioral differences and the computational processes underlying them.

### Amygdala involvement of older adults is lower in trust decisions, but higher in outcome evaluation

Each trial in our repeated trust game task consisted of four phases (Figure 3a), the preparation phase where participants were presented with the trustworthy or untrustworthy trustee’s face, the investment phase to execute their trust decisions, a waiting phase for the trustee’s feedback, and finally the outcome phase where the trustee’s back-transfer was revealed. In previous work (on the younger part of the present sample(Sladky et al., 2021)), we had demonstrated the importance of analyzing these task phases separately. This had shown that the central (CeA) and basolateral amygdala (BLA) as well as the basal forebrain (BF) are crucial for successful trust learning, corroborating the findings of a human amygdala lesion study (Rosenberger et al., 2019). Other areas such as the nucleus accumbens (NAc), bed nucleus of the stria terminalis (BST), and substantia nigra/ ventral tegmental area (SN+VTA), were related to reward feedback processing in the outcome phase. To extend these findings to the putative effects of aging and loneliness on trust learning, we applied the same fMRI preprocessing and analysis strategy used in our previous publication to ensure consistency.

**Figure 3:**
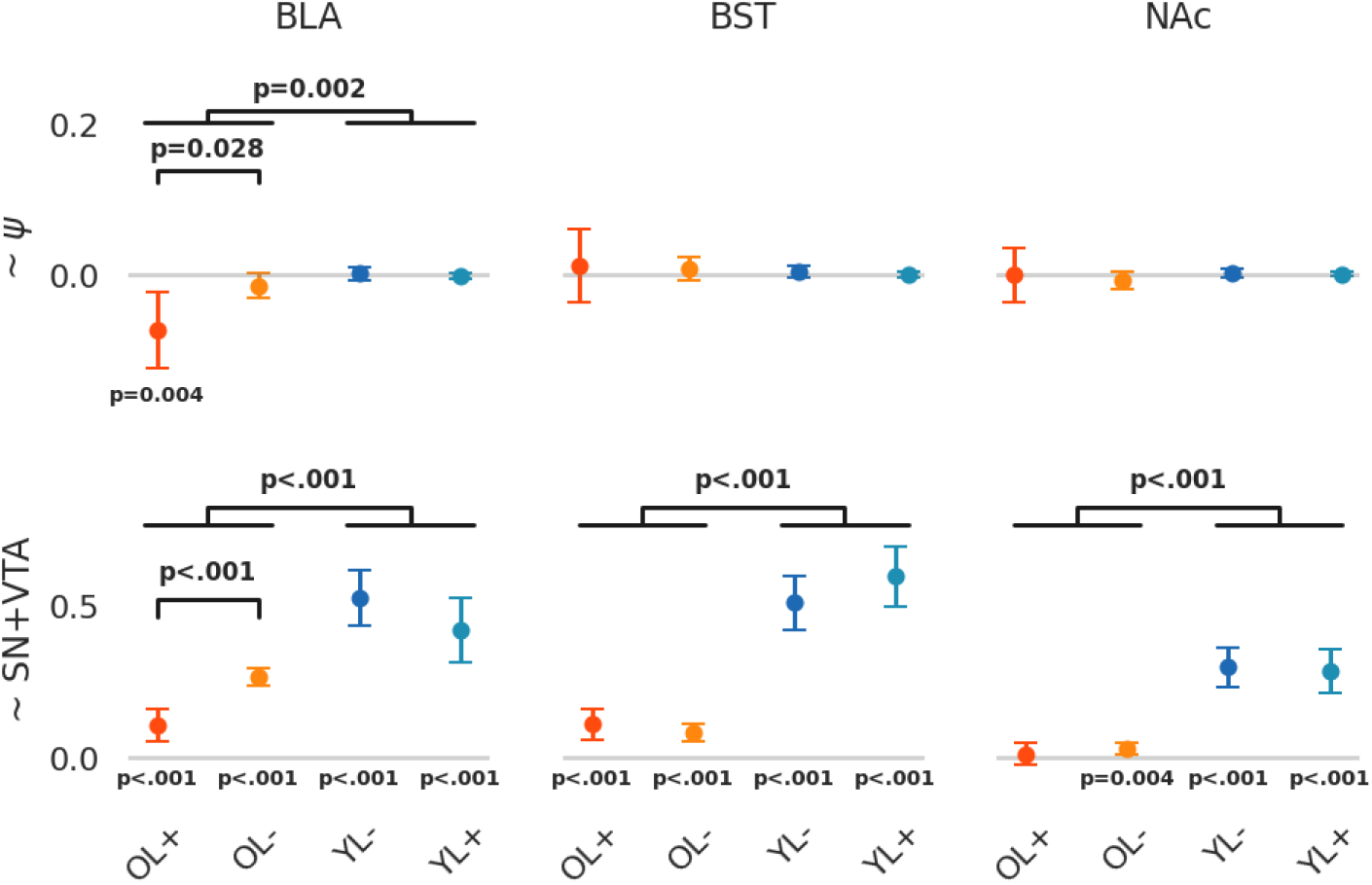
Coupling between VOI activation during trust learning (i.e., the outcome event when the back-transfer feedback is presented) and the learning rate (ψ) and SN+VTA activation. BLA, BST, and NAc responses were positively coupled with SN+VTA activation but to a lesser degree in older adults. Lower BLA response is also associated with a lower learning rate (t(2983)=−2.00, p=0.045) and when differentiating for groups, there were significant age differences (Y>O: p=0.002) additionally increased by loneliness (OL−>OL+: p=0.004).

Consequently, we first investigated how age and loneliness influence activation in the amygdala subregions (BLA, CeA, BST) during the preparation phase. In the preparation phase, amygdala (BLA, CeA, BST) activations were lower in older adults (significant two-sample t-tests of β estimates from the HRF model, Figure 3b). More specifically, during preparation we observed more activation in younger adults in the BLA (F(1,110)=8.76, p=0.004, η^2^(partial)=0.02), the CeA (F(1,110)=8.19, p=0.005, η^2^(partial)=0.03), and also the BST (F(1,110)=6.93, p=0.010, η^2^(partial)=2.93e-03). Loneliness main and interaction effects were not significant (all p>0.068).

However, for the outcome event (the beginning of the outcome phase), older adults showed more amygdala activation. We found significant age effects in the BLA (F(1,110)=11.94, p<.001, η^2^(partial)=7.96e-03) and the CeA (F(1,110)=14.36, p<.001, η^2^(partial)=0.09), where we also observed significant effects for loneliness (F(1,110)=11.04, p=0.001, η^2^(partial)=0.04) and the interaction age × loneliness (F(1,110)=4.17, p=0.043, η^2^(partial)=0.12) (Figure 3c). This means, higher amygdala activation does not automatically entail better belief updating, which was lower in older adults. These findings were suggestive of the coupling between trial-by-trial learning rate and dopaminergic signaling to play a role, which we targeted in our next set of analyses. As dynamic trust learning was predicted to not only involve the amygdala but also the nucleus accumbens (NAc, relevant for action selection and reward processing primarily via dopaminergic mechanisms), the cholinergic basal forebrain (BF, assumed to regulate the amygdala during trust decisions and learning), and the substantia nigra and ventral tegmental area (SN+VTA, whose dopaminergic input is assumed to regulate precision in the amygdala and NAc), we focused on these areas.

For NAc, during preparation, we observed statistically significant effects for age (F(1,110)=19.20, p<.001, η^2^(partial)=0.01) and the interaction with loneliness (F(1,110)=3.95, p=0.049, η^2^(partial)=0.15). Post-hoc t-tests showed higher NAc activation in younger adults (Y>O: p<0.001) interacting with loneliness (F(1,110)=3.95, p=0.049, η^2^(partial)=0.15), with a difference in non-lonely but not in lonely individuals (YL−>OL−: p<0.001, YL+>OL+: p=0.365). During outcome evaluation, we observed statistically significant effects for age (Y>O: F(1,110)=14.77, p<.001, η^2^(partial)=0.06), loneliness (L->L+) F(1,110)=7.03, p=0.009, η^2^(partial)=0.07) and the interaction (F(1,110)=7.85, p=0.006, η^2^(partial)=0.12). Here, NAc activation was lower in Y vs. O and within the O group lonely participants had lower NAc activity (OL−>OL+: p=0.045) (Figure 3c).

For BF, during preparation, we also observed statistically significant effects for age (Y>O: F(1,110)=5.87, p=0.017, η^2^(partial)=6.28e-03) but not for loneliness. Significant effects were also seen for the outcome event for age (O>Y: F(1,110)=11.56, p<.001, η^2^(partial)=0.03) but no effects for loneliness. We thus assume that cholinergic signaling is lower during trust behavior preparation but higher during reward feedback processing at older age (Figure 3c).

For SN+VTA, during preparation, we also observed effects for age (Y>O: F(1,110)=15.08, p<.001, η^2^(partial)=0.02) but not for loneliness. For the outcome event we observed no significant age-effect (p=660), however for loneliness (L->L+: F(1,110)=10.40, p=0.002, η^2^(partial)=0.06) and the interaction (F(1,110)=7.32, p=0.008, η^2^(partial)=1.76e-03). Post-hoc t-tests showed that while there are no differences between YL− vs. OL− (p=0.971), OL− showed higher SN+VTA activation than OL+ (p=0.009) (Figure 3c).

### Lower learning in older adults – particularly those who feel lonely

The results of the computational modeling analyses suggest that older adults, particularly those who are lonely, have a lower learning rate (ψ, trial-by-trial precision-weight on the prediction error) and more inflexible beliefs (ω, volatility beliefs), both affecting learning (ψ, ω are higher in learners). The brain analyses suggest that during the outcome phase, activation in the amygdala (BLA, CeA) as well as BF, NAc is higher in older adults – however, lower in the dopaminergic SN+VTA in lonely older adults. This prompted us to explore how, during belief updating, trial-by-trail ψ related to neural activity in the outcome phase in the task-relevant areas.

As a first step, we validated if and how ψ is related to BOLD responses during the outcome phase. This revealed a negative relationship (VOI ~ ψ + (1 | Subject)) in the BLA (t(2983)=−2.00, p=0.045), the BST (t(2763)=−2.02, p=0.044), and the NAc (t(3180)=−4.92, p<0.001), suggesting that with higher activity in these regions during outcome evaluation, learning from feedback is impaired. CeA, BF, and SN+VTA did not show significant relationships with the learning rate (p>0.219).

Then, we calculated an LMM (VOI ~ ψ × SN+VTA × Group + Trustee + (1 | Subject)) testing for possible coupling between the VOI activation (BLA, BST, and NAc) with SN+VTA and learning rate. This was confirmed with model comparison against the simpler model without SN+VTA (all p<0.001). We used the for subgroups as grouping variable (OL+,OL−,YL−,YL+) and tested the hypotheses of a higher coupling in younger people (Y>O) and that loneliness additionally reduces the coupling in older adults (OL−>OL+).

The coupling between ψ and the BLA was indeed higher in younger adults (Y>O: p=0.002) and also higher in non-lonely older adults (OL−>OL+: p=0.28, Figure 4). The coupling between SN+VTA and all VOIs was positive (p<0.001) but lower in older adults (Y>O: p<0.001) and in the BLA even lower in lonely older adults (OL−>OL+: p<0.001, Figure 4).

Taken together, these analyses suggest that the interactions between amygdala and NAc are relevant for trust learning in the different subgroups. Of interest, even though activation in these areas is consistently higher in older adults (Figure 3c), the coupling with the SN+VTA activation (as a proxy for dopaminergic signaling) is lower, particularly if they are also more lonely. This suggests that reduced dopaminergic signaling could impair reward signaling or belief updating in the BLA.

## Discussion

In this study, we investigated how age and loneliness affect trust and trust learning. We found marked age-related differences in the behavioral, neuro-computational, and neuroimaging results, which are in some aspects aggravated by loneliness. Older adults displayed both lower initial trust, lower overall trust, and lower trust learning. In real life, this may translate to higher skepticism towards trust partners, and disadvantages in both social interactions and financial investments. Of note, these effects differ for trustworthy vs. untrustworthy interaction partners. The finding that older adults are more trusting than younger adults towards the untrustworthy trustee (who displayed negative indicators of trustworthiness) align well with the meta-analysis by (Bailey and Leon, 2019), who identified a moderate age-related increase in trust in response to neutral and negative indicators of trustworthiness. For the initial trust behavior, i.e., investments that are not informed by the trustee’s observed back-transfer behavior, older adults generally invested less than younger adults. One explanation for this lower baseline trust could be that older adults avoid risks in social interactions. This may either be the result of their more extensive negative learning history, or of a general tendency towards more cautious behavior with a lower tendency for risky decision-making (Deakin et al., 2004; Mamerow et al., 2016), which may serve as an optimal compensatory behavior if they are (implicitly) aware of their deficits in trust learning.

Our data reveal the strongest difference in learning rates between lonely older adults and younger adults. Future studies should determine if (a) older people are more susceptible to loneliness-effects or (b) the impaired trust learning in older adults could be the consequence of long-term experience of chronic loneliness, resulting in entrenched beliefs about other people and loss of trust learning abilities. It has been suggested that older adults rely more on preserved knowledge (Hedden and Gabrieli, 2004), which would be in line with the notion that the brain ages optimally to model the (social) environment (Moran et al., 2014) given the metabolic constraints imposed by aging (Shaulson et al., 2024). However, a downside of increased cognitive and metabolic efficiency, evinced by reliance on heuristics and prior knowledge, is reduced flexibility, manifesting as impaired learning from the consequences of one’s actions.

Impaired learning could be caused by reduced sensitivity to feedback in general (tied to reward and reward sensitivity) or, more specifically, to the integration of prediction errors (resulting in differences in the updating of trustworthiness beliefs). Older adults responded stronger to trust behavior outcomes in the BLA, CeA, NAc, regions known to be involved in the encoding of a variety of different types of prediction error (Corlett et al., 2022). While this suggests a higher feedback sensitivity on the one hand, it also indicated that the feedback and the resulting prediction errors are not as effectively utilized as learning signals.

There is growing consensus that blunted or less effective learning could be the consequence of reduced dopamine signaling, as suggested by recent reviews (e.g., (Orsini et al., 2015)) on the role of dopamine in regulating PFC-subcortical networks during risky decision-making tasks or specifically in the context of trust learning (Schuster and Lamm, 2025). Moreover, we had previously shown that blocking D2/3 receptors, which increases available dopamine through D1 receptors, enhances belief volatility and increases precision weights on prediction errors (Mikus et al., 2023). Using the same task and model, the younger participants of our study replicate the findings from the placebo group in Mikus et al.(Mikus et al., 2023), also confirming that beliefs about untrustworthy individuals appear more volatile (Mikus et al., 2023; Siegel et al., 2018). The older adults in our study, however, exhibit significantly lower volatility and precision weights, consistent with the idea of reduced dopamine availability and, consequently, less effective updating of beliefs and therefore suboptimal outcomes for trust behavior. This aligns also with the finding that lonely older adults indeed showed less activity and involvement of the dopaminergic midbrain in the assumed BLA-based belief update mechanism.

Age-related declines in dopaminergic function have been well-documented: there are decreases of dopamine receptors, midbrain dopamine transporter expression, dopamine transporter availability, and D2 receptor density, which is associated with lower glucose metabolism in regions like the frontal cortex, anterior cingulate, temporal cortex, and caudate nucleus (Bannon and Whitty, 1997; Rinne et al., 1990); also see (Hedden and Gabrieli, 2004) for review. Furthermore, fMRI studies indicate that aging impairs reward-based learning and decision-making, while, on the other hand, dopamine therapy can improve learning from rewarding outcomes in older adults (Chowdhury et al., 2013; Samanez-Larkin and Knutson, 2015). Even a single dose of L-dopa was found to support lasting fear extinction learning (Esser et al., 2021; Haaker et al., 2013), which could also be useful for unlearning avoidance-driven distrust. Moreover, our own prior research using a similar task design showed that increasing available dopamine by using the D2/3-receptor blocker Sulpiride increased trust learning rates (Mikus et al., 2023). Despite these lines of evidence suggest that dopaminergic medication could improve cognitive function (Mehta and Riedel, 2006) and, thus, social cognition in older adults, there are two important caveats. First, a memory encoding study showed that Sulpiride compared to the D2-like agonist Bromocriptine had opposite effects on age-related memory impairments (Morcom et al., 2010), highlighting the lifetime plasticity of receptor coupling mechanisms and their behavioral consequences. Second, pharmacological unspecific upregulation of dopamine is also not a panacea (Kulisevsky, 2000). While the D2/3-receptor blocker Haloperidol can increase recognition accuracy in a memory task and at the same time increase activation in the amygdala, SN/VTA and other areas, which would be desirable for trust learning, it was also found to decrease the subjective confidence in one’s decisions (Clos et al., 2019). However, this alteration in one’s metacognitive abilities could also be one of the mechanisms that contributes to improved learning by relaxing top-down beliefs, e.g. about harmful intent attributions as shown in a dictator/sharing game after Haloperidol administration (Barnby et al., 2024), which corresponds to the broader idea of an inverted-U shape of dopamine availability for optimal cognitive functions (Cools and D’Esposito, 2011) allowing an adaptive balance of belief updating and protection (Schuster and Lamm, 2025).

Beyond the effects of aging, the additional role that loneliness had in our study also need consideration. Loneliness does not only affect social cognition but also various brain functions and based on different levels and neurotransmitter systems (for review see Cacioppo and Hawkley, 2009; Vitale and Smith, 2022). In humans, reward-related ventral striatum activation was lower when lonely individuals viewed images of unfamiliar people compared to their non-lonely counterparts (Cacioppo et al., 2009). In contrast, lonely individuals showed heightened activity in the ventral striatum when viewing familiar faces versus strangers (Inagaki et al., 2016). Interestingly, while lonely individuals display dampened activity at rest and in response to various positive social and non-social stimuli, non-lonely individuals exhibit higher activation in the insula, anterior cingulate cortex, prefrontal cortex, and ventral striatum when they feel excluded in a virtual ball game (Eisenberger et al., 2003). These observations all point towards a role of dopaminergic reward and reward anticipation related processes, linking loneliness and trust in similar ways as aging and trust. Indeed, it has even been shown that acute social isolation results in craving-like responses, as similar midbrain responses are observed in individuals who have fasted towards food-related stimuli in the nucleus accumbens and socially excluded participants towards social stimuli in the caudate nucleus (Stijovic et al., 2023; Tomova et al., 2020). While the precise mechanisms are still unclear, rodent data suggests that while VTA dopaminergic signaling corresponds to social reward, dopaminergic neurons in the dorsal raphe nucleus could alter this effect due to synaptic changes following acute social isolation (Matthews et al., 2016). The interaction effect we found for the age and loneliness interaction could also support the assumed opposite roles of acute and chronic isolation (Matthews and Tye, 2019): while acute isolation promotes pro-social motivation, chronic isolation has the opposite effect.

Beyond the neural mechanisms, our study bears wider implications for healthy aging. While some recent accounts suggest a higher baseline trust in older people (Bailey and Leon, 2019), our results suggest lower initial trust. Also, despite the popular trope, there is no evidence that older adults are disproportionately more susceptible to consumer fraud (Ross et al., 2014); however, our results suggest that there is less effective learning from negative outcomes in older people and financial exploitation risk could be particularly high for people with dementia (Wong et al., 2017). How our experimental findings translate to real-life decision-making and long-term behaviors needs further research and validation, though. Impaired trust learning may also result in missed opportunities for positive social interactions, implying negative health consequences beyond psycho-social wellbeing. How social factors can impair health has been shown even in non-human studies. E.g., social stress shortens the lifespan via higher risk of cardiovascular disease in mice (Razzoli et al., 2018) or inflammations due to altered immune regulation/response in macaques (Snyder-Mackler et al., 2016) and other species (reviewed in Snyder-Mackler et al., 2020). Loss of social participation, leading to loneliness, can result from social stigma associated with aging (Pillemer et al., 2021) but also multi-facetted reasons from individual somatic causes, e.g., age-related hearing loss (Brewster et al., 2021), to causes affecting the social habitat, e.g., suboptimal urban planning (Belsky and Baccarelli, 2023). Yet, changing these factors is thought to delay aging effects (Wang et al., 2021).

## Conclusion

Our findings reveal that impaired social cognition may stem from brain aging and loneliness, driven by reduced dopaminergic efficacy, which can make it difficult for affected individuals to form new social connections. As loneliness accelerates brain aging and disrupts dopaminergic function, this research underscores a potentially reversible vicious cycle that warrants further investigation and targeted psycho-social interventions to mitigate loneliness in older adults.

## Materials and Methods

### Participants

In total 114 participants participated in our study: 52 healthy, neurotypical older adults (64 to 84 years) and 62 younger adults (age: 20 to 33) (Demographical data are reported in **Error! Reference source not found.)**.

Exclusion criteria were standard MRI exclusion criteria (e.g.: pregnancy, claustrophobia, and MRI-incompatible implants, clinically significant somatic diseases), a history of psychiatric or neurological disorders, substance abuse, psychopharmacological medication, less than nine years of education, as well as not being task-naive (e.g., having already participated in a similar study or being a psychology student). All participants provided written informed consent in accordance with the Declaration of Helsinki and were compensated for their participation. The study was approved by the ethics committee of the Medical University of Vienna (EK-Nr. 1489/2015).

### Participant characteristics

A chi-squared test revealed no difference in the number of participants (χ^2^(1, N=114)=0.0003, p=0.987) and their gender (χ^2^(3, N=114)=1.608, p=0.6576) across the groups. In our sample, younger adults had more years of formal education, with a medium effect size (ANOVA: Education ~ Age Group × Loneliness Group, F(1,110)=13.09, p<.001; η2 (partial)=0.11, 95% CI [0.03, 1.00]). The main effect of Loneliness group (p=0.290) and the interaction Age × Loneliness group (p=0.991) were not significant.

By design, self-reported loneliness was higher with a large effect size in the high Loneliness Group (L+>L-, ANOVA Loneliness Scale ~ Age Group × Loneliness Group, F(1, 110)=210.84, p<.001; η2 (partial)=0.66, 95% CI [0.57, 1.00]). In addition, the main effect of Age Group (Y>O) is statistically significant and medium (F(1,110)=14.68, p<.001, η2 (partial)=0.12) and the interaction between Age Group and Loneliness Group is statistically significant and small (F(1, 110)=6.04, p=0.016; η2 (partial)=0.05, 95% CI [5.48e-03, 1.00]). The post-hoc t-test revealed no difference between the L-groups (p=0.7917532) but the younger adults in the L+ group were more lonely than the L+ older adults (mean difference=6.91, out of scale values between 20 to 80, 95% CI [2.87, 10.96], p<0.001).

In sum, while the sample was balanced in terms of group size, age range, and gender distribution, younger adults reported more years of formal education and loneliness scores were higher in the younger high-loneliness Group (YL+) compared to the older high-loneliness Group (OL+).

### Procedure and Task

This study was part of a bigger project including two additional tasks, which are not reported in the current article (Riva et al., 2023, 2022). The task and procedure were identical in our previously published study using data of the younger sample (Sladky et al., 2024); the data and analyses reported here are independent of the previous report with respect to (a) not being biased by results or procedures developed on the younger adult sample (e.g. because they were tailored to the characteristics of the younger adults sample and thus could favor that sample with respect to hypotheses related to age-related differences) (b) being novel and not presented elsewhere, as the present report includes the older adults group and data on loneliness of the participants. Moreover, the previous report did not use a computational model to study the processes underlying trust behavior and trust learning. Participants were initially invited to a screening session where they performed cognitive tasks and completed self-reported measures of psychological traits. The main session typically took place within two weeks of the screening session. Upon arrival at the MRI facility (University of Vienna MR Center), participants were accompanied by two other individuals, who in reality were two confederates of the experimenter, playing the roles of the trustees. After signing the consent form and completing the MR safety questionnaire, participants and confederates were introduced to the entire session protocol. Subsequently, they underwent training on the three tasks, including the trust game. At the end of the training, participants were required to answer questions to ensure their understanding of the task. Finally, participants were placed in the MRI scanner, while the confederates were positioned in the computer room adjacent to the scanner room, ostensibly playing the task.

The repeated trust game was adapted from our previous study (Rosenberger et al., 2019) and programmed in z-Tree (version 3.3.7). The script of this trust game is deposited online (Rosenberger et al., 2019) and already described in (Sladky et al., 2021): Two players, an investor (participant) and one of the trustees (confederate), engaged in a monetary exchange game to maximize their outcomes. At the start of each round (20 rounds per trustworthy condition and 20 rounds per untrustworthy condition), the investor received an endowment of 10 monetary units to invest in the trustee (1 to 10 monetary units). The confederates, who were allegedly played by the participants in an alternate randomized order, took on the trustee roles and back-transferred more or less of the investment. However, their actions were preprogrammed to ensure one confederate behaved trustworthily while the other acted untrustworthily. The confederates/players were of similar age and gender to the participant. Each round encompasses four phases. In the *preparation* phase, participants are presented with the picture of the trustee’s face they are playing with in the current round. In the *investment* phase, participants invest (part of) their endowment; the investment is tripled and then transferred to the trustee. During the *waiting* phase, the trustee ostensibly performs their back-transfers. Finally, during the *outcome* phase, participants are presented with the back-transfer outcome. In the first two rounds, both the trustworthy and untrustworthy trustee back-transferred the same amount of the money invested to the participants. In the following rounds, the trustworthy trustee always back-transferred as much or more than the money invested by the participant, whereas the untrustworthy trustee always back-transferred less than or as much as the money invested by the participant. The sums invested by the participants were considered as a measure of trust given to the two trustees and used as the main variable of interest. Points earned throughout the task were transformed to Euros and added to the participants’ compensation.

At the end of the task, participants were presented with the trustees’ picture and were asked to rate them on four adjectives: trustworthiness, fairness, attractiveness, and intelligence (original German: *Wie vertrauenswürdig/fair/attraktiv/intelligent haben Sie den/die Teilnehmer/in wahrgenommen?*). Ratings were provided on visual analogue scales and transformed off-line to a numerical range between −10 and +10.

### Behavioral data analysis

It is commonly understood that participants’ investment behavior is a behavioral expression of how they judged the players’ trustworthiness and changes reflect the extent to which they updated their beliefs (Bellucci et al., 2017; Chang et al., 2010; Rosenberger et al., 2019; Sladky et al., 2021). This *objective* measure of trust was used to distinguish between learners and non-learners using the median as cut-off value from (Sladky et al., 2021).

The hierarchical Gaussian filter (HGF) computational modeling was performed as described in (Mikus et al., 2023) using the original R-code provided in https://github.com/nacemikus/belief-volatility-da-trustgame.git.

In essence, based on the participant’s behavior, we employ the HGF to model the subjective trustworthiness beliefs associated with each of the two trustees. These trustworthiness beliefs are represented as Gaussians N(μ_t_, σ_t_), initially set to μ_0_, an estimator of the participant’s uninformed beliefs. Over time, these beliefs are allowed to evolve through precision-weighted belief updates, which serve as a dynamic learning rate (ψ_t_) governed by prediction error and confidence in prior beliefs and likelihood. Precision, in this context, is inversely proportional to variance. An additional hierarchical level is used to account for higher-level volatility beliefs. These belief volatility parameters ω for the trustworthy and untrustworthy trustee govern the rate of change of σ_t_via the precision-weights ψ_t_ that serve as dynamic learning rates. Identical to the original publication, we estimate initial trustworthiness belief per participant (μ_0_). In other words, we estimate the latent belief about the trustee’s trustworthiness as a Gaussian distribution with a specific mean (representing how trustworthy I think the trustee is) and variance (indicating my confidence in my assessment). Higher belief volatility (the amount of variance I can expect in the trustee’s trustworthiness) implies higher variance (or lower precision), suggesting an unpredictable trustee. The resulting dynamic learning rate (ψ_t_) on the prediction error is proportional to the expected variance or inversely proportional to the precision of beliefs and is therefore referred to as a precision-weight. Higher precision-weighted learning rates and low precision of prior beliefs evince stronger changes in beliefs. The ordinal logistic link function governs how beliefs about others’ trustworthiness map to investments with two additional subject-level parameters: choice uncertainty (γ) and the slope (η) (**Error! Reference source not found**. a). The parameter estimation is done through hierarchical Bayesian inference, where we estimate all individual and group-level parameters in one inferential step. Model comparison against different versions of the HGF and Rescorla-Wagner models was performed in the original publication of the current model (Mikus et al., 2023) and the present work uses the same winning model. The mathematical formalism is described in detail in the original HGF publication (Mathys et al., 2014) and the Methods section of the current model (Mikus et al., 2023).

### Functional MRI Data acquisition, processing, and analyses

As described in (Sladky et al., 2021), MRI acquisitions were performed on a Skyra 3 Tesla MRI scanner (Siemens Healthineers, Erlangen, Germany) using the manufacturer’s 32 channel head coil at the MR Center of the University of Vienna. In a single session, one run of the repeated trust game was performed by the participant while we performed functional MRI using a gradient echo T2*-weighted echo planar image sequence with the following parameters: MB-EPI factor=4, TR/TE=704/34 ms, 2.2×2.2×3.5 mm^3^, 96×92×32 voxels, flip angle=50°, n<2400 volumes.

Data processing and analyses of the functional MRI data were performed in SPM (SPM12, http://www.fil.ion.ucl.ac.uk/spm/software/spm12/) and the Python packages nipype (http://nipy.org/nipype) and nilearn (http://nilearn.github.io). Preprocessing comprised slice-timing correction (Sladky et al., 2011) using SPM, realignment using SPM, non-linear normalization of the EPI images (Calhoun et al., 2017)to a study-specific group template using ANTs (Avants et al., 2011), and spatial smoothing with a 6 mm FWHM Gaussian kernel using SPM. SPM result maps were warped from study space to MNI space (final resolution=1.5LJ×LJ1.5LJ×LJ1.5LJmm^3^) using ANTs. VOI analyses were performed in study space using anatomical masks that were transformed to the study-specific group template using the inverse transformation from MNI using ANTs’ *ApplyTransforms*. To verify that the EPI images properly covered the VOI, axial slices of the median single-subject mean volumes in study space are presented in the supplementary material of our previous publication (Sladky et al., 2021, Supplementary Figure 1). An additional functional connectivity analysis was performed to verify that the specificity of the BOLD signal is sufficient for distinguishing between the BLA and CeA. To this end, task fMRI data were corrected for white matter and CSF signal and task effects (Ganger et al., 2015) using regression before estimation of the functional connectivity maps of the BLA and CeA seeds (Sladky et al., 2021, Supplementary Figure 2).

First-level analyses of the data were implemented using nipype and performed using SPM12’s GLM approach. The GLM design matrix encompassed individual regressors for each of the 4 task phases (i.e., preparation, investment, waiting, and outcome) and each of the 2 interaction partners (trustworthy and untrustworthy, resulting in 8 effects of interest. Additionally, 6 realignment parameters were added as nuisance regressors to account for residual head motion effects. Second-level analyses of the data were implemented using nilearn’s group-level approach.

Volume of interest analyses were performed on the mean timeseries extracted using nilearn’s *fit_transform* from anatomical masks from the BLA, CeA (Tyszka and Pauli, 2016), NAc (AAL Atlas), BST (Torrisi et al., 2015), SN/VTA (Talairach atlas transformed to MNI space), and basal forebrain (Jülich Brain MPM atlas). To investigate phase-dependent activation in the amygdala subregions, timeseries analyses were conducted based on the estimated percent signal change, using custom python scripts that reproduced SPM’s default GLM analysis, using SPM’s canonical HRF to convolve the 8 regressors of interest (4 phases × 2 players), using the realignment parameters as confounds and a DCT-based high-pass filter with SPM’s default *f*=1/128 Hz cut-off frequency to account for signal drifts.

For consistency with the previous fMRI publication (Sladky et al., 2021), frequentist statistical analyses of fMRI data was performed. Parameter estimates were converted to percent signal change. Group effects (older vs. younger adults) were compared using t-tests using Python scipy and reproduced in R, yielding the same results. Analyses of the multivariate effects (age, loneliness; age, loneliness, trustee; age, loneliness, learners; loneliness, learners) were performed in R using car::Anova.

## Supplemental Content

### Supplementary Results

#### Behavioral Data

As a measure for the amount of learning we also investigated changes in absolute trust behavior across the rounds (Figure 1e) and particularly focused on the difference between the last two rounds, as indicator for uncertainty because trust behavior change should decrease over time. An ANOVA (Investment Change ~ Age × Loneliness) revealed significant main effects for Age Group (O>Y, F(1,110)=11.94, p<.001, η^2^(partial)=0.10) and interaction effects for age and loneliness (F(1,110)=4.59, p=0.034, η^2^(partial)=0.04). The post-hoc t-tests were significant for OL+>YL− (p=0.027) and OL+>YL+ (p<0.001). This means, with accumulating evidence about the two trustees, younger adults showed less changes in their investment behavior, indicating that they have made up their mind (or learned) about the trustees’ trustworthiness and that this impacted their behavior. In older adults, particularly in those who were lonely, the investment changes did not reduce as much, indicating less learning or smaller effects of learning on behavior.

Finally, connecting trust behavior to subjective trustworthiness beliefs using the subjective trustworthiness rating difference delivered at the end of all 20 rounds, we found a significant effect for age (Y>O, F(1,103)=10.02, p=0.002, η^2^(partial)=0.01) but not for loneliness or the interaction (all p>0.156); note that rating data from seven older adults were not successfully recorded and thus not included in this analysis (Figure 1a).

**Figure S1:**
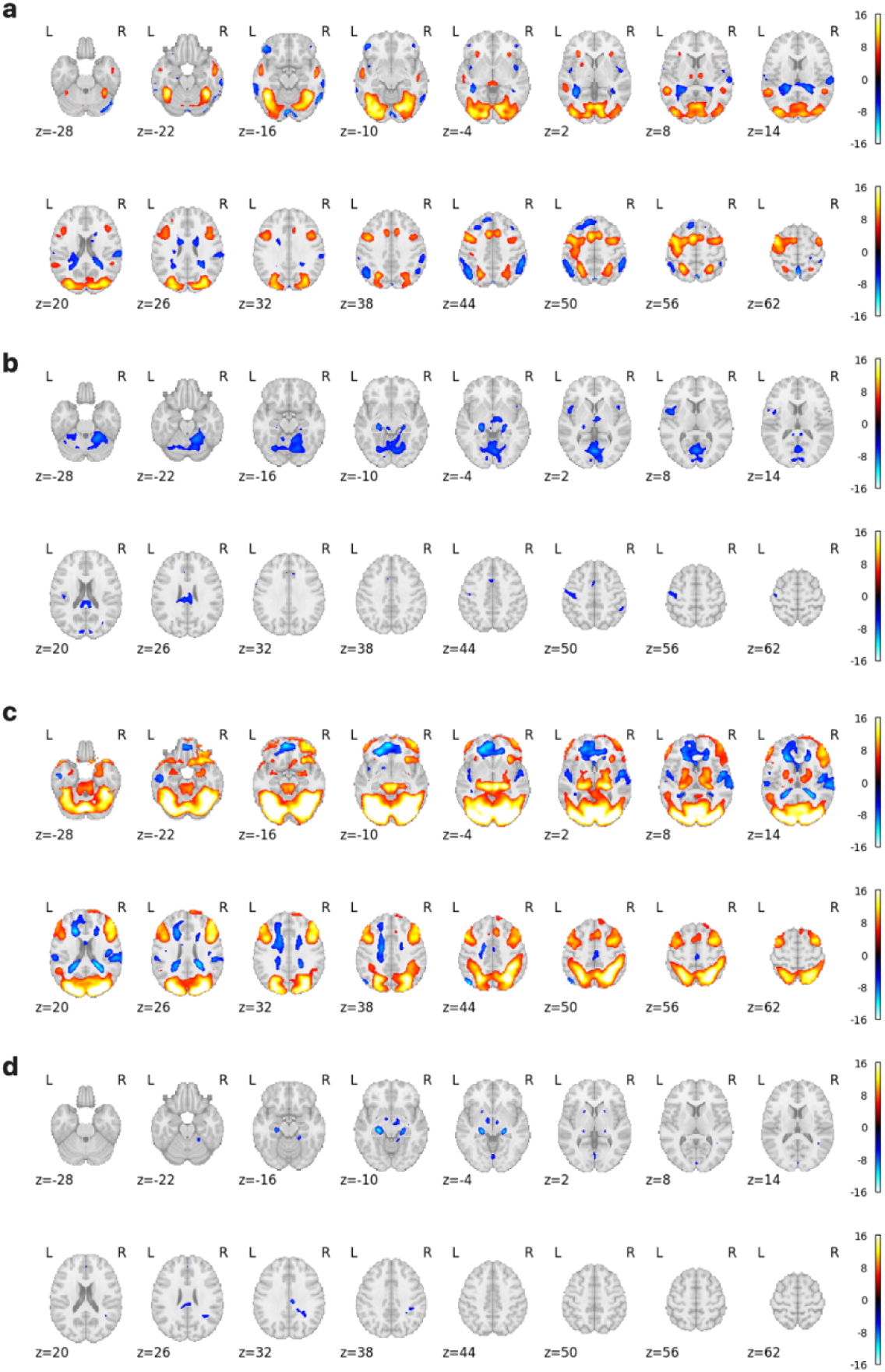
Whole brain fMRI results. **a.** Preparation phase. Activaton in visual areas and thalamus, temporal lobe, anterior insula, and (dorsolateral) prefrontal cortex are consistent with demanding stochastic visual-cogntive decision making. **b.** Age effects during preparation phase. In addition to the hypothesis-driven findings, we observed age-related hypOctivity in the interoception-related anterior insula and memory-related posterior hippocampus, which could contribute to the lower efficiency of trust behavior in older adults. **c.** Outcome phase. Activation was comparable to the preparation phase but stronger, particularly in the amygdala region, posterior hippocampus, midbrain, and dorsal striatum consistent with reward processing. Dorsomedial prefrontal cortex was hypOctive. **d.** Age effects during outcome phase. Most prominent age-related hypOctivity were observed in the posterior hippocampus. All p<0.05 FWE whole brain corrected. Loneliness or Age*Loneliness interaction were not significant using strict thresholding.

#### Functional MRI

For completeness, we also present the whole brain analysis results in Figure S1. We used functional connectivity as an additional validation that the specificity of our data is sufficient to discriminate the amygdala subnuclei, which we presented in the supplement of our previous publication (Sladky et al., 2021).

Of note, the shift from amygdala hypoactivation in the preparation to hyperactivation in the outcome phase in older adults can already be observed during the investment and waiting periods in the BLA (Figure 3). This means that this transition may not be related to outcome evaluations specifically but could be related to reward anticipation that should take place between having made a trust decision and the revelation of the decision by the trustee. However, this finding regarding activation dynamics across task phases should be considered exploratory. It is not directly motivated by the results from our previous studies (Rosenberger et al., 2019; Sladky et al., 2021) and its interpretation remains preliminary, considering that our task design may not be sensitive enough to distinguish alternative explanations, such as a delayed or prolonged trust decision influencing the BOLD response after the preparation phase without affecting responsiveness during outcome evaluation.

#### HGF Model

To investigate what factors could affect the trial-by-trial learning rate (ψ), we calculated a series of linear mixed effects models.

First, we wanted to investigate whether the learning rate actually captures the expected learning goal to differentiate between the trustees (LRN+/− learners group definition based on the median split in the total investment differences 14) and if it is influenced by the trustee (T+/− depending on trustworthiness) (ψ ~ Learners Group × Trustee + (1 | subject)). We found that ψ is higher in learners (LRN+ vs. LRN-, t(131)=22.29) and for the untrustworthy trustee (T− vs. T+, t(4216)=−10.64) with a significant interaction (t(4216)=−39.97), all p<0.001). This is a face validation of ψ as a computational marker for learning. Second, for the aging and loneliness effects (ψ ~ Trustee × Age Group × Loneliness Group + (1 | subject)) all main effects and interactions were significant (p<0.001) (Figure 2c). ψ was higher for the untrustworthy trustee, younger adults, and loneliness amplified this difference. For the trustworthy trustee there were no such effects.

We investigated the full model ψ ~ Learners Group × Age Group × Loneliness Group + Trustee + (1 | subject) and found a significant positive effect for LRN+ (t(106)=4.19, p<0.001), the untrustworthy trustee T− (t(4217)=34.08, p<0.001). Most importantly, there was an interaction between LRN+ and L+ (t(106)=3.06, p=0.003) and LRN+ and L+ and O (t(106)=−2.30, p=0.023): In LRN+ we observed no difference between YL− and OL−, however, in YL+ ψ was higher compared to OL+ (Figure 2d).

## Statistics and Reproducibility

In total 114 participants participated in our study: 52 healthy, neurotypical older adults (64 to 84 years) and 62 younger adults (age: 20 to 33) (Demographical data are reported in **Error! Reference source not found.**). Data were processed and analyzed using SPM (SPM12, http://www.fil.ion.ucl.ac.uk/spm/software/spm12/) and the Python projects nipype (http://nipy.org/nipype) and nilearn (http://nilearn.github.io). Additional statistical analyses were performed in R and Python using numpy and scipy; for visualization matplotlib and nilearn were used. Code and data required to reproduce our results and figures are publicly available on https://github.com/scanunit.

## Data availability

The data needed to reproduce the results and figures are published on our lab’s public github page (https://github.com/scanunit).

## Code availability

The code needed to reproduce the results and figures are published on our lab’s public github page (https://github.com/scanunit).

## Acknowledgments

We thank Helena Hartmann for her help in collecting the data. This work was funded by the Austrian Science Fund FWF (P29150, and in part by 10.55776/COE16). For open access purposes, the author has applied a CC BY public copyright license to any author accepted manuscript version arising from this submission. Claus Lamm acknowledges partial funding from the Vienna Science and Technology Fund (WWTF VRG13-007).

## Supplementary Information

Supplementary Information can be found online at [tbc].

## Author Contributions

Conceptualization and Methodology, R.S., F.R., C.L.; Investigation, F.R.; Formal Analysis, R.S.; Writing – Original Draft, R.S., F.R., C.L.; Writing – Review & Editing, R.S., F.R., C.L.; Funding Acquisition, C.L.

